# Activation of the cGAS-STING pathway contributes to cancer-related fatigue in a murine model of head and neck cancer

**DOI:** 10.1101/2025.10.02.680158

**Authors:** Brandon Chelette, Abate Bashaw, Joshua D. Bryant, Kiersten Scott, Chinenye Chidomere, A. Phillip West, Robert Dantzer

**Affiliations:** Department of Symptom Research, The University of Texas MD Anderson Cancer Center, Houston, TX 77030, USA; The Jackson Laboratory, Bar Harbor, ME 04609, USA; Department of Microbial Pathogenesis and Immunology, Texas A&M University College of Medicine, Bryan, TX 77807, USA

**Author notes:** Corresponding Authors: A. Phillip West, PhD, The Jackson Laboratory, Bar Harbor, ME 04609, Robert Dantzer, DVM, PhD, Department of Symptom Research, The University of Texas MD Anderson Cancer Center, Houston, TX 77030. These authors contributed equally (Brandon Chelette, Abate Bashaw).

## Abstract

Cancer-related inflammation and metabolic alterations extend beyond the tumor microenvironment, exerting systemic effects that disrupt energy homeostasis and contribute to reduced physical function and chronic fatigue. The cGAS–STING pathway has emerged as a key regulator of innate immunity and inflammation; however, its role in cancer-associated fatigue remains poorly understood. In this study, we investigated the contribution of cGAS– STING–mediated inflammation to cancer- and/or its treatment-induced fatigue using a mouse model of human papillomavirus-related head and neck cancer. Wheel running activity, along with inflammatory and metabolic changes in tumor and liver tissues, were assessed following chemoradiotherapy and pharmacological inhibition of STING in tumor-bearing and tumor-free control mice. The results revealed that tumor growth and chemoradiotherapy activated the cGAS-STING pathway, together with an upregulation of proinflammatory mediators and alterations of mitochondrial and metabolic gene expression in the liver. To inhibit STING activation, mice were administered H-151, a specific STING antagonist. This intervention attenuated hepatic inflammatory signatures and mitigated tumor and/or chemoradiotherapy-associated behavioral fatigue measured by decreased voluntary wheel running. These findings implicate for the first time the cGAS–STING signaling pathway and metabolic homeostasis in cancer- and cancer therapy-related fatigue.

## 1. Introduction

Cancer-related fatigue is a distressing, persistent feeling of physical, emotional, and/or cognitive exhaustion related to cancer or cancer treatment, which significantly impacts the daily functions of patients (1-5). It affects approximately 20% of patients at the time of diagnosis, increases in incidence and severity during cancer therapy, and persists in about 30% of patients for months and even years after treatment ends. Fatigue develops as a non-specific symptom of innate immune activation in several disease conditions, largely due to the ability of peripheral inflammation to propagate to the brain through multiple immune-to-brain communication pathways (6). Chronic inflammation and carcinogenesis are closely interconnected, and as cancer patients exhibit high levels of inflammatory mediators in the circulation, systemic inflammation is considered a major culprit in cancer-related fatigue (7). However, the inflammatory response associated with cancer differs from that triggered by pathogen-associated molecular patterns, both in its origin and in its cellular and molecular characteristics;□.

Cancer is a dynamic and complex disease characterized by intricate interactions between tumor cells, immune cells, and stromal cells of the tumor microenvironment. Cancer-associated inflammation has both intrinsic and extrinsic components. Tumor cell intrinsic inflammation is triggered by the mechanisms involved in cell transformation and proliferation, including oncogenes, genomic instability, and metabolic reprogramming to respond to the increased requirements of highly proliferative tumor cells in energy and building blocks. Extrinsic inflammation originates in the tissue that surrounds the tumor in response either to damage caused by the expanding tumor or to secondary infection, and it is mediated by infiltrating innate immune cells. For example, macrophages in the tumor microenvironment can be co-opted by the tumor to serve its metabolic needs (8). Mitochondria are crucial in both recognizing cellular stress and producing damage-associated molecular patterns (DAMPs) (9), which are key factors in initiating immune responses and inflammation (8). Genomic instability and oxidative stress, common features of cancer, cause damage and release of nuclear and mitochondrial DNA into the cytosol and extracellular space, where they can activate innate immune responses and inflammation. The cyclic GMP-AMP synthase–stimulator of interferon genes (cGAS-STING) pathway is a central regulator of double stranded DNA sensing and type I interferon signaling, playing a critical role in mediating the immune response to DNA released during cellular stress or damage. Cancer therapies add another layer of genomic stress. Many cancer therapies, including chemotherapy and radiotherapy, induce mitochondrial and nuclear DNA damage, leading to the release of DNA from both cancerous and healthy cells, thereby triggering inflammation. Dysregulated activation of cGAS-STING pathway has been implicated in various inflammatory conditions and may contribute to treatment-associated inflammation and toxicity (10-12).

In addition to tumor cell-intrinsic effects, cancer cells actively remodel the microenvironment of distant organs to support the demands of rapid tumor proliferation. They recruit metabolically active organs such as skeletal muscles and the liver, disrupting systemic metabolic homeostasis. This cancer metabolism syndrome has been primarily studied in the context of cancer cachexia^1^□, resulting in impaired mitochondrial metabolism that leads to ineffective adenosine triphosphate (ATP) generation, lipid alterations in the mitochondria, and increased expression of uncoupling proteins. Moreover, mounting evidence suggests that mitochondrial dysfunction caused by cancer and/or cancer treatment alters neuronal functions, contributing to the development of chemotherapy-induced peripheral neuropathies and cognitive dysfunction(13-16). However, the role of mitochondrial dysfunction in the initiation and persistence of cancer-related fatigue is not yet fully understood.

As the cGAS-STING pathway is strongly activated by both cancer and cancer therapy, the present series of experiments was initiated to assess the role of this pathway in cancer-related fatigue. We employed a syngeneic mouse model of human papillomavirus (HPV)-associated head and neck squamous cell carcinoma (HNSCC) derived from mouse oropharyngeal epithelial cells that stably express HPV-16 E6 and E7 genes along with H-Ras oncogene (mEER)(17, 18) to investigate the role of cancer- and chemoradiotherapy (CRT)-associated cGAS-STING activation in cancer-related fatigue. E6 and E7 inhibit apoptosis by inactivating p53 and pRB pathways, while E6 synergizes with H-Ras to promote proliferation (17, 18). The advantage of the mEER model is that it generates highly metabolically active tumor (19) but in a context of relative immunosuppression, as E6 and E7 counteract the activation of both nuclear factor kappa B (NF-κB) and the cGAS-STING pathway. However, this is partly compensated by H-Ras overexpression as it promotes proliferation and inflammation by activating the transcription factors STAT3 and C/EBPb (20). Therefore, using this model allows us to study the effects of CRT on mitochondrial function and immune activation in the periphery without introducing confounding immune effects from the tumors. We show that tumor growth and CRT induce distinct transcriptional changes in the liver, characterized by activation of the cGAS-STING pathway and disruption of mitochondrial and metabolic homeostasis. Importantly, we reveal that treatment with the STING antagonist H-151 (21) attenuates hepatic inflammatory signatures and improves voluntary wheel-running activity, indicating a critical role for the cGAS-STING pathway in cancer- and CRT-associated fatigue.

## 2. Materials and Methods

### 2.1. Animals

Male C57BL/6J mice (at 10 weeks age) were obtained from The Jackson Laboratory and maintained in a temperature and humidity-controlled room on a 12 h light-dark cycle with food and water provided ad libitum. Mice were individually housed at the start of each experiment. All animal experiments were approved by the Institutional Animal Care and Use Committee (IACUC) of the University of Texas MD Anderson Cancer Center and carried out in accordance with the NIH Guide for the Care and Use of Laboratory Animals. Male mice were exclusively used in the study because sex differences in the intensity and duration of our primary outcome, behavioral fatigue, measured by decreases in voluntary wheel running, are not apparent in the development of fatgue but only its recovery (22), which was not studied here.

### 2.2. In vivo tumor growth and therapeutics

mEER tumor transplantation was carried out as previously described (22). Briefly, 12-week-old male mice were subcutaneously injected with 1 × 10^6^ mEER cells suspended in 100 ml of sterile phosphate-buffered saline (PBS). Control mice received vehicle PBS. Tumor cells were injected either in the right flank of mice that did not receive chemoradiotherapy or into the right hind leg to allow local irradiation of the tumor without impacting internal organs. Tumor volume was measured using digital calipers in three orthogonal dimensions, and volume was calculated by the formula (volume = (π/6) x (d1 x d2 x d3). Tumor weight was used as an alternative measure of tumor growth in some experiments where the tumor mass was more diffuse, impairing accurate caliper measurements. Mice were monitored daily for general health and euthanized earlier than initially scheduled if their tumor diameter was superior or equal to 15 mm, or if they displayed lethargy, lost more than 10% body weight, or showed other signs of profound sickness or distress. Body weight was measured on a regular basis during the duration of the experiment.

Cisplatin chemotherapy combined with radiotherapy (chemoradiotherapy, CRT) was used to treat tumors weekly, beginning 7 days after injection of mEER cells. Cisplatin was administered intraperitoneally at a dose of 5.28 mg/kg. Radiotherapy was administered at the dose of 8 Gy using a cesium irradiator (22).

We initially used mice with a whole-body genetic deletion of STING to assess the role of the cGAS-STING pathway in cancer-related fatigue (B6(Cg)-*Tme m173*^*tm1*.*2Camb*^/J, Strain #025805, Jackson Lab). However, we found that these mice were deficient in several behavioral aspects, including voluntary wheel running and cognition. There are also reports of hypersensitivity to nociceptive stimuli and signs of hyperalgesia (23). Therefore, we switched to a pharmacological approach and selected H-151, a highly potent and selective small molecule inhibitor of STING. H-151 covalently targets STING’s transmembrane cysteine residue, thereby blocking its activation-induced palmitoylation (21). H-151 was administered either to mEER tumor-bearing mice to assess the role of the cGAS-STING pathway in the fatigue associated with tumor growth, or to tumor-bearing mice treated with CRT to determine the role of this pathway in the behavioral fatigue induced by cancer therapy. In both cases, H-151 (Invivogen, inh-h151) was diluted in DMSO and further diluted in 10% Tween in PBS, before being administered intraperitoneally at a dose of 750 nmol or 210 mg per mouse.

### 2.3. Voluntary wheel running activity

Mice were provided access to running wheels placed into their home cage as previously described (24). After stabilization of their running activity, which usually required 2 weeks, the number of revolutions during the night was automatically recorded and expressed as a percent of their baseline activity before interventions (implantation of tumor cells with or without CRT).

### 2.4. Tissue collection

A separate experiment using a 2 (± mEER tumor cells) x 2 (± 3 weekly cycles of CRT beginning 7 days after implantation of mEER tumor cells) with n=7 male mice per group was carried out to assess the effects of the tumor and its treatment on the cGAS-STING pathway. Mice were euthanized by CO_2_ inhalation and perfused intracardially with PBS. Euthanasia took place 24 h after the last cycle of CRT. Tissues (brains, livers, and tumors) were collected and snap frozen in liquid nitrogen and stored at□−80□°C for qRT-PCR transcript analyses and western blot protein analysis.

### 2.5. Quantitative reverse transcription PCR using custom PCR array

Total RNA was isolated from snap-frozen liver samples using RNA-Solv Reagent (Omega Bio-Tek; Norcross, GA, USA) following the manufacturer’s protocol. RNA was quantified, and approximately 1□µg of total RNA per sample was reverse transcribed into cDNA using the RT^2^ First Strand Kit (Qiagen) A custom PCR array for genes related to inflammation, interferon signaling, metabolism, and mitochondria dynamics (Qiagen) was performed using RT^2^ SYBR Green qPCR Mastermix (Qiagen) and a CFX384 Real-Time PCR Detection System (BioRad). The genes were selected based on our previous studies on the effect of mEER tumors and CRT on the host (25). All samples were tested in duplicate. Eukaryotic Translation Initiation Factor 3 Subunit A (*Eif3a*) and Ribosomal Protein S3 (*Rps3*) were used as internal reference genes. Relative expression was calculated using the ΔΔCt method, with normalization to reference genes. Fold changes were computed relative to control, where positive values indicate upregulation and negative values indicate downregulation.

### 2.6. Immunoblotting

Protein lysates were prepared from frozen samples using 1% NP-40 lysis buffer supplemented with a protease inhibitors (ThermoFisher Scientific). Total protein concentrations were determined by Micro BCA Protein Assay kit (Thermo Scientific). Equal amounts of protein from each sample were separated by acrylamide gel electrophoresis and transferred to polyvinylidene fluoride (PVDF) membranes (Bio-Rad). After air drying, membranes were incubated with primary antibodies overnight at 4°C with gentle agitation. The following primary antibodies were used: STING (D2P2F), STAT1, Phospho-p65 (93H1), RIG-I (4200), NLRP3 (D4D8T), and IκB-ζ (93726) from Cell Signaling Technology, ZBP1 (aa 1-411, AdipoGen), P49 (IFIT3, Ganes Sen, Cleveland Clinic), GAPDH (60004-1-Ig, Proteintech), and HSP60 (Santa Cruz Biotechnology). Membranes were washed and incubated with appropriate horseradish peroxidase-conjugated secondary antibodies for 1 hour at room temperature. Protein bands were developed using an Immobilon Crescendo Western HRP Substrate and visualized via chemiluminescence imaging or X-ray film. The relative protein abundance was quantified using ImageJ software, with normalization performed against loading controls. Statistical analysis was done by GraphPad Prism and are presented as □ ± □standard error of the mean.

### 2.7. Statistical analysis

All data are presented as mean ± standard error of the mean and analyzed by two-way or three-way ANOVA with repeated measurements on the time factor for *in vivo* longitudinal studies. Post hoc comparisons were conducted using the Tukey HSD test for multiple comparisons. P values equal to or less than 0.05 were considered statistically significant but trend values were also reported for the qRT-PCR results.

## 3. Results

### 3.1. Cancer and its treatment activate the cGAS-STING pathway

Cancer-related inflammation and metabolic alterations extend beyond tumor-localized effects, exerting systemic impacts that disrupt multiple organ systems and contribute to impaired energy homeostasis and reduced physical function. To examine hepatic inflammation and metabolic alterations in response to cancer and its treatment, we performed mRNA expression analysis using a custom qRT-PCR array in mEER tumor bearing or tumor free control mice, with or without CRT. A 2 × 2 factorial design was employed to assess the independent and interactive effects of tumor and CRT. The results revealed that tumor growth and CRT induce distinct transcriptional changes in the liver, characterized by activation of the cGAS-STING pathway and disruption of mitochondrial and metabolic homeostasis (**Fig. 1**). Notably, upregulated expression of *Mb21b1* (encoding cGAS), *Tmem173* (encoding STING), and several interferon-stimulated genes (ISGs) were detected in livers of tumor-bearing mice compared to tumor-free mice, indicating activation of the cGAS-STING pathway (**Fig. 1**). *Itgam* (encoding CD11b; a marker of myeloid cells) and proinflammatory genes, particularly *Il1b* and *Ccl2*, were also elevated in the livers of tumor-bearing mice. Interestingly, CRT reduced the expression of most of the cGAS-STING pathway-related genes observed in untreated tumor-bearing mice. It also partially attenuated the expression of *Il1b* and *Itgam* (**Fig.1**). This reduction in the inflammatory signature may reflect transcriptomic reprogramming toward immunosuppressive pathways in response to repeated CRT.

**Fig. 1.**
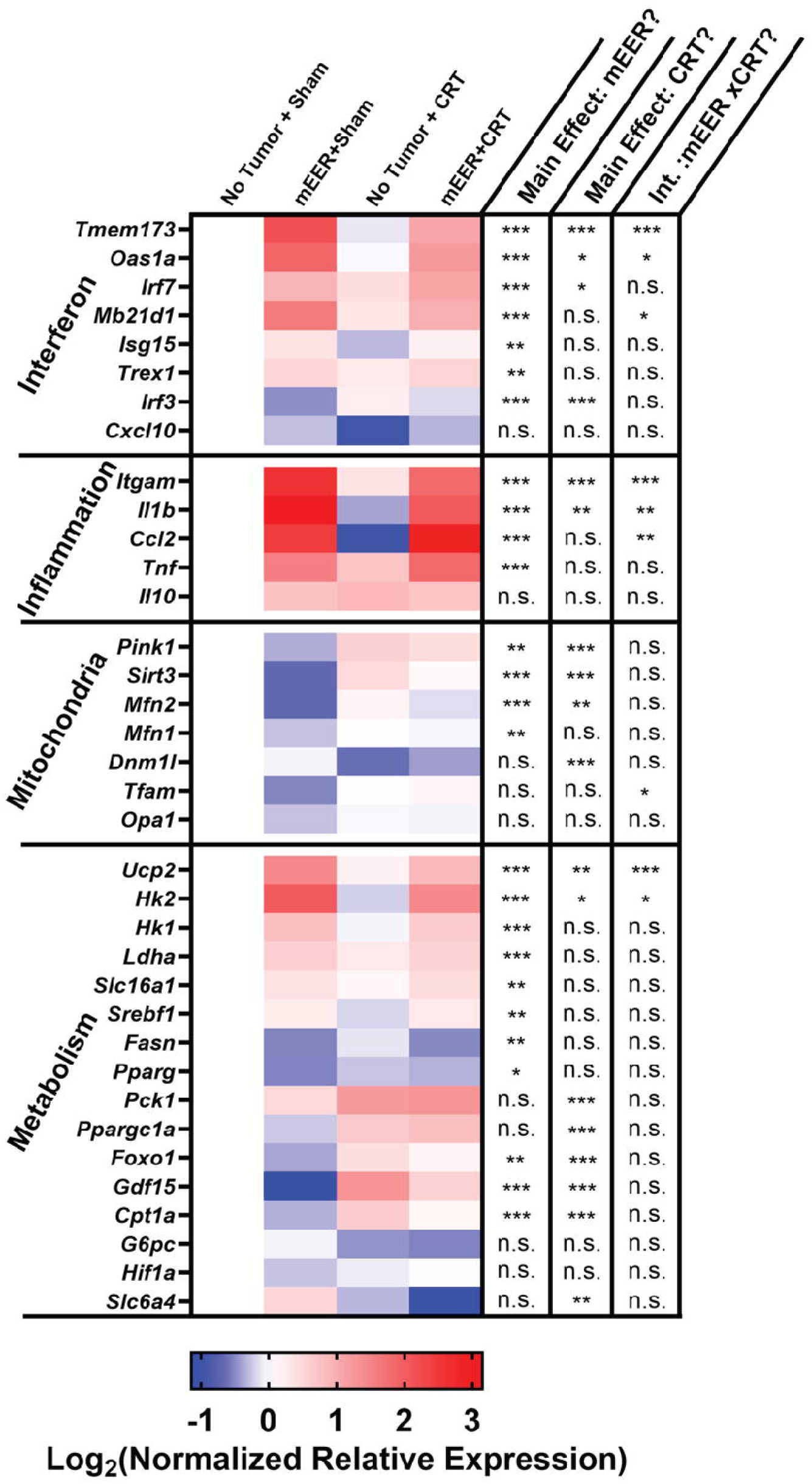
mEER tumor growth and chemoradiotherapy significantly altered the liver transcriptome. Heatmap showing the effects of mEER tumor and/or chemoradiotherapy (CRT) on the expression of genes related to interferon signaling, inflammation, metabolism, and mitochondrial dynamics. A custom quantitative reverse-transcription polymerase chain reaction was used to measure gene expression levels in the liver tissues. The color gradient highlights differential expression patterns in response to the tumor and/or CRT. Red indicates upregulation, while blue represents downregulation, relative to untreated tumor-free mice. * p<0.05, **p<0.01, *** p<0.001, NS not significant (2-way ANOVA).

A significant alteration in the transcriptomic signatures for liver metabolic and mitochondrial function was also observed in response to tumor growth and CRT. Downregulated expression of genes involved in mitochondrial quality control, including *Pink1, Sirt3, Mfn1*, and *Mfn2*, was detected in the livers of tumor-bearing mice (**Fig. 1**). Interestingly, CRT increased most of these genes compared to sham-treated controls. Other metabolic genes such as *Ucp2, Hk1, Hk2*, and *Srebf1* were significantly upregulated in tumor-bearing mice, whereas *Pck1, Cpt1a, Foxo1, Ppargc1a* (encoding PGC-1α), and *Gdf15* were downregulated. CRT induced a significant increase of these downregulated genes in both tumor-bearing and tumor-free mice. The upregulation of *Hk1* and *Hk2* in tumor bearing mice, which encode the glycolytic enzymes hexokinase 1 and 2, along with *Srebf1* that regulates lipogenesis, suggests enhanced glycolytic activity and lipid synthesis while downregulation of *Ppargc1a* (PGC-1α), the key regulator of mitochondrial biogenesis, indicates impaired mitochondrial function. These data reflect tumor associated systemic inflammation and impaired mitochondrial metabolism that favor glycolysis and lipogenesis.

Consistent with the gene expression results, we observed significantly higher protein levels of inflammatory molecules in liver lysates isolated from mEER-tumor-bearing mice in comparison with tumor-free control mice (**Fig. 2 A,C**). The cytosolic innate immune sensors ZBP1, RIG-I, and NLRP3, and downstream signaling molecule STING were significantly increased in the livers of mEER-tumor-bearing mice (**Fig. 2 A,C**). Notably, CRT in non-tumor-bearing mice did not impact inflammatory protein expression in the liver. CRT also did not significantly alter innate immune activation in the livers of mEER-bearing-mice, although there was a modest decrease in some proteins. In contrast, tumor samples from CRT-treated mice exhibited increased RIG-I and STAT1 levels, indicating the potential link between CRT-induced tumor death, damage-associated molecular pattern (DAMP) release, and subsequent innate immune signaling (**Fig. 2B, D**). These data confirm activation of the cGAS-STING pathway and inflammatory signaling at the biochemical level in tumor-bearing mice and indicate a close relationship between inflammation and alterations in mitochondrial dynamics and energy metabolism in the liver in response to cancer and its treatment.

**Fig. 2.**
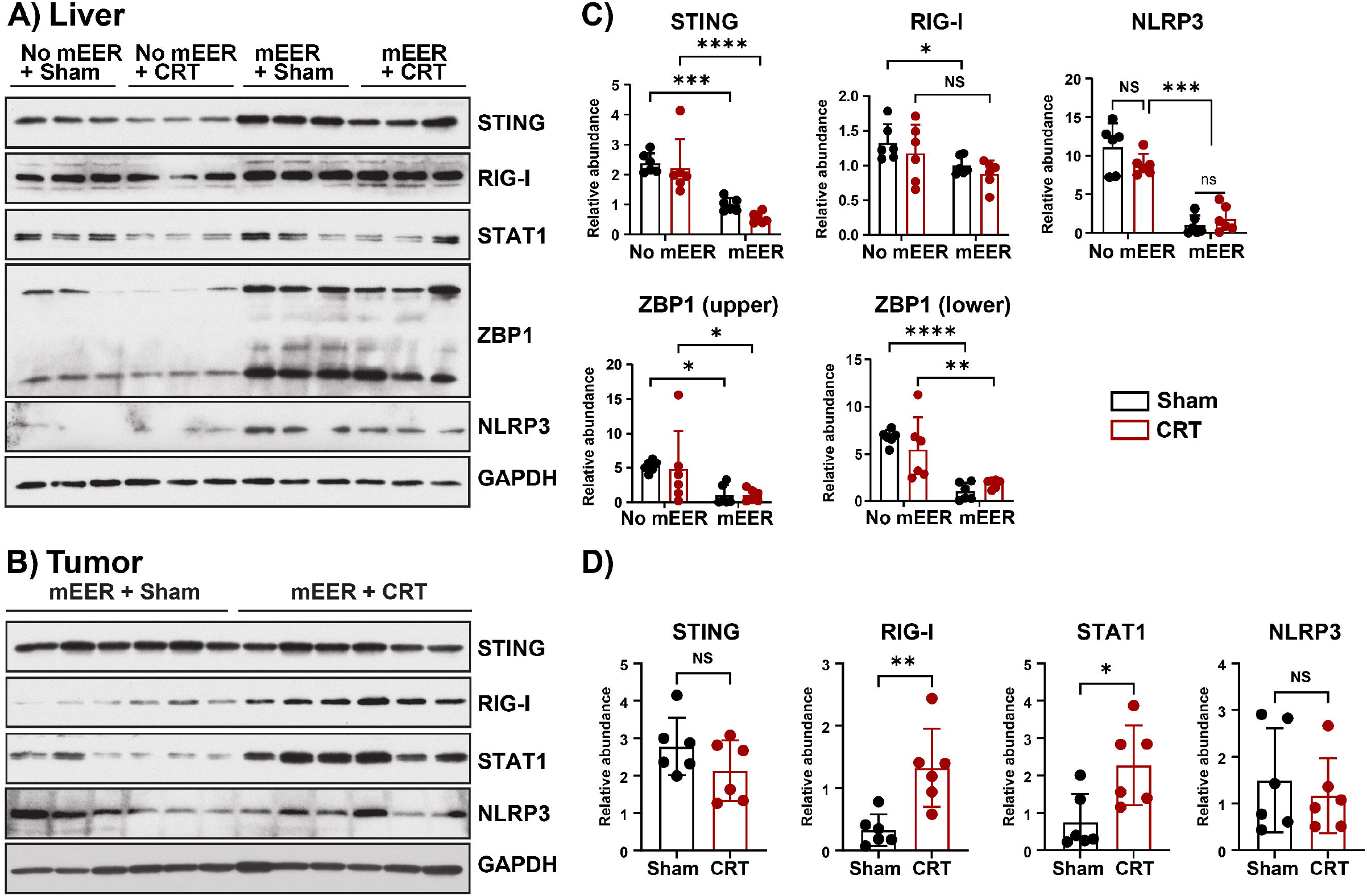
Western blot analysis of inflammatory response-related proteins in liver or tumor lysates from mEER tumor-bearing or control mice. Representative western blots of indicated proteins in liver (A) and tumor samples (B) from mEER tumor-bearing and control mice, with or without chemoradiotherapy (CRT). (C) Bar plots depicting relative protein abundance in liver tissue lysates from mEER tumor-bearing and control mice treated with CRT or untreated controls (sham). (D) Protein levels in tumor samples from mEER mice treated with CRT or untreated controls. Densitometry quantification of each western blot band was analyzed using ImageJ software and normalized to GAPDH loading control. Comparisons in (C) one-way ANOVA, multiple comparisons test, and in (D) unpaired *t*-test, two-tailed were performed using GraphPad Prism. *p < 0.05, **p < 0.01, ***p < 0.001, ****p < 0.0001, ns: not significant. Error bars, SEM.

### 3.2. Targeting STING with H-151 attenuates tumor and CRT-induced decrease in voluntary wheel running and activation of liver inflammation

To next assess whether inhibition of the STING–mediated inflammatory response can alleviate cancer- and cancer therapy-related fatigue, STING antagonist H-151 was administered in two independent experiments carried out successively on tumor-bearing mice (Experiment I) and on tumor-bearing mice submitted to CRT (Experiment II).

#### 3.2.1. Experiment I

In this experiment, mEER tumor-bearing mice were treated with H-151 or PBS vehicle every other day, starting 10 days after tumor transplantation and continuing until the end of the study. H-151 was prepared as described in the methods section and injected intraperitoneally at a dose of 750 nmol. A 2 × 2 factorial design was used to assess the effects of tumor burden (± mEER) and STING inhibition (± H-151), with 7–8 male mice per group (**Fig. 3A**).

**Fig. 3:**
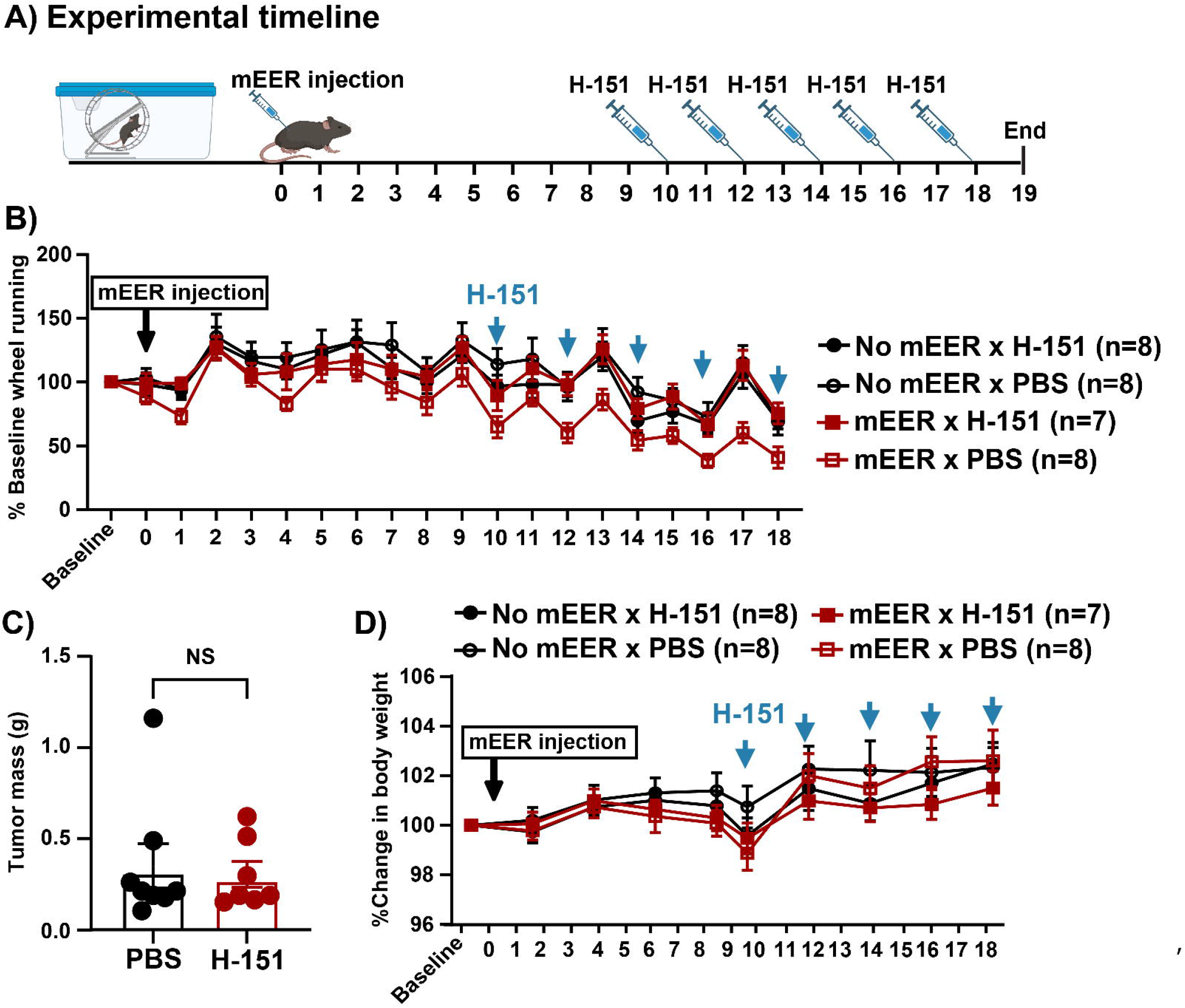
STING antagonism by administration of H-151 attenuates the development of behavioral fatigue measured by decreased wheel running in mEER tumor-bearing mice. (A) Schematic of the experimental timeline. (B) Variations over time of nightly running wheel counts expressed as percent of baseline in mice implanted or not with mEER tumor cells and treated or not with H-151 (mean +/-SEM, n=7-8 mice per group). (C) Tumor mass measured at the end of the experiment in tumor-bearing mice treated or not with H-151. (D) Variations over time of body weight expressed as a percentage of baseline in the same mice as in (A).

Wheel running performance varied over time (**Fig. 3B**), with a general tendency to decrease by the end of the experiment as it is often the case in mice that are wheel running for an extended period of time (a little longer than one month in the present experiment) (time factor F(11, 297)=46.1, p<0.001). Compared to vehicle-treated control mice, each injection of H-151 treatment improved wheel running time of the mEER tumor-bearing mice (treatment factor F(1, 27)=5.09, p<0.05; tumor x treatment interaction F(1, 27)=4.49, p<0.05). H-151 treatment had no significant effect on tumor size measured at the end of the experiment (t = 0.32, ns; Fig. 3C). Body weight increased over time (time effect: F (5,135) = 23.4, *p* < 0.001); however, this increase was more pronounced in mEER tumor-bearing mice that received the vehicle, as indicated by a significant tumor × H-151 × time interaction (F (5, 135) = 2.36, *p* < 0.05; Fig. 3D).

The biochemical assays carried out on the livers and tumors revealed an increased expression of STING, as well as ZBP1 and IFIT3-interferon-inducible genes (ISGs) downstream of the cGAS-STING signaling pathway in the liver lysates of tumor-bearing mice compared to tumor-free control mice (**Fig. 4 A,C)**. These elevated levels were reversed by H-151 treatment, confirming that tumor growth activates the cGAS-STING pathway in the liver and mediates tumor-related stress-induced inflammatory responses. STING antagonist had no significant effect on the levels of the ISGs analyzed in mEER tumors (**Fig. 4 B,D**), suggesting that the effects of H-151 on wheel running performance are mediated by impacting peripheral inflammation and metabolic rewiring.

**Fig. 4.**
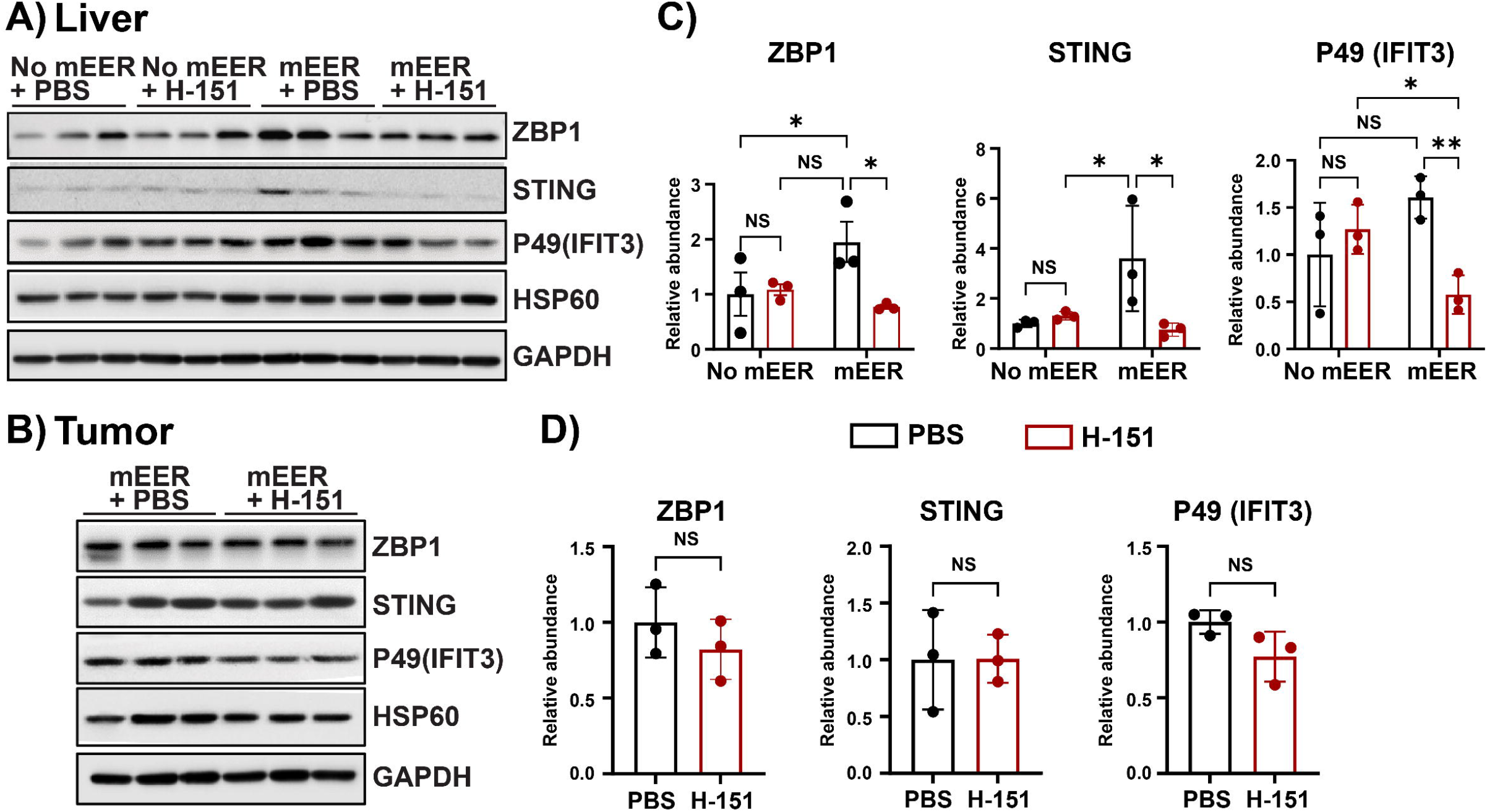
H-151 STING inhibitor reduced activation of liver interferon and inflammatory factors in mEER tumor-bearing mice. Representative Western blots of liver (A) and tumor (B) samples from mEER tumor-bearing and control mice treated with H-151 or PBS control. (C) Relative protein levels in liver samples from H-151- or vehicle (PBS)-treated mice. (D) Indicated protein levels in tumor samples from H-151- or vehicle (PBS)-treated mice. Densitometry quantification of each western blot band was analyzed using ImageJ software and normalized to HSP60 and GAPDH loading controls. Comparisons in (C) one-way ANOVA, multiple comparisons test, and in (D) unpaired *t*-test, two-tailed were performed using GraphPad Prism. *p < 0.05, **p < 0.01, ***p < 0.001, ****p < 0.0001, ns: not significant. Error bars = SEM.

#### 3.2.2. Experiment II

This experiment was carried out to further investigate whether the STING antagonist H-151 mitigates CRT-asociated fatigue in tumor-bearing mice. Mice bearing a mEER tumor were administered H-151 twice, on the day before CRT and on the day of CRT. This was repeated for each cycle of CRT using a 2 (± mEER/CRT) x 2 (± H-151) factorial design with n=7 - 8 male mice per group (**Fig. 5A**). Control mice were not implanted with mEER tumor cells and did not receive CRT. A decrease in wheel running activity was observed following the transplantation of the mEER tumor cells, with a further decrease after each CRT session (**Fig. 5B**). This decrease was followed by a recovery, which was initially apparent after the first CRT session but became blunted after the second session. The tumor + CRT effect was highly significant (F(1, 27) =29.4, p<0.001), with the intensity of this effect varying over time (tumor x time F (15, 405) =2.72, p=0.001). The H-151 effect was not significant (F(1, 27) =0.28), but it interacted significantly with time (H-151 x time, F(15, 405=1.89, p<0.05). There was a trend for interaction between the tumor + CRT factor and the H-151 factor (F (1, 27) =2.92, p<0.10), indicating that the effect of H-151 tended to be larger in tumor-bearing mice that received CRT than in control mice. The interaction between the tumor + CRT, H-151, and time did not reach significance (F (15, 405)=1.42). To analyze the effects of H-151 on the decrease in wheel running induced by CRT in tumor-bearing mice more specifically, a separate ANOVA with repeated measurements on the time factor was performed on the data collected during H-151 administration and CRT (from Day 6 to Day 21 of the experiment). The results of this analysis confirmed those already obtained, with a significant effect of the time (F(15, 390)=13.8, p<0.001) and tumor + CRT factors (F(1, 26)=54.3, p<0.001) and the H-151 x tumor + CRT interaction (F(1, 26)=8.32, p<0.01). Like before, the interaction between the tumor + CRT, H-151, and time did not reach significance (F(15, 390)=1.54).

**Fig. 5.**
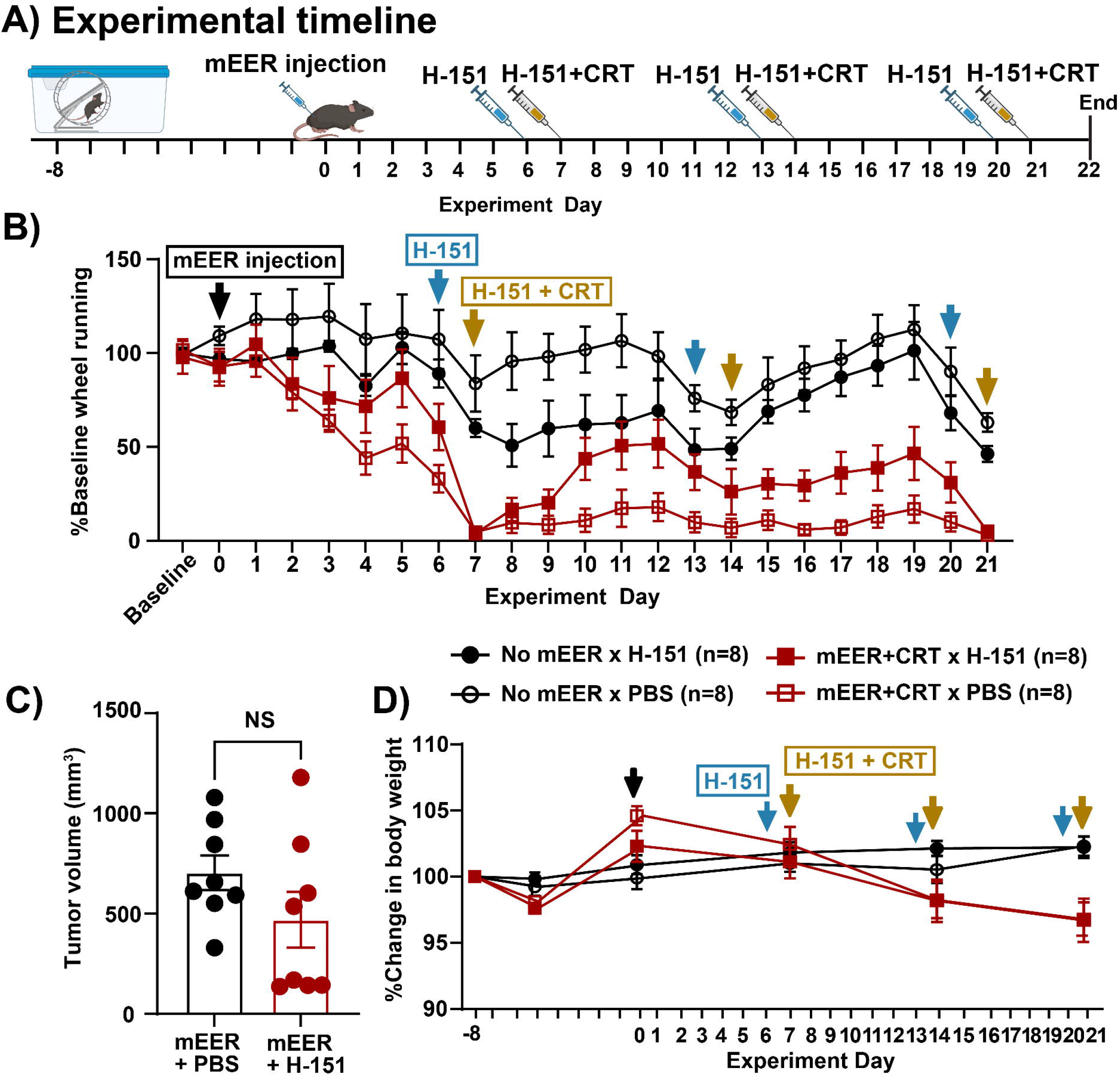
STING antagonism by administration of H-151 attenuates the development of behavioral fatigue measured by decreased wheel running in mEER tumor-bearing mice treated or not by chemoradiotherapy (CRT). (A) Schematic of the experimental timeline. (B) Variations over time of nightly running wheel counts expressed as a percent of baseline in mice implanted or not with mEER tumor cells and exposed to three weekly cycles of CRT beginning 7 days after implantation of mEER tumor cells and treated or not with H-151 (mean +/-SEM, n=7-8 mice per group). (C) Tumor mass measured at the end of the experiment in tumor-bearing mice administered CRT and treated or not with H-151. (D) Variations over time of body weight expressed as a percentage of baseline in the same mice as in A.

These data indicate that the H-151-treated tumor-bearing mice were less impaired in their wheel running during CRT. The administration of H-151 did not affect tumor volume measured at the end of the experiment (**Fig. 5C**). Body weight expressed as a percentage of baseline decreased during the last two sessions of CRT independently of H-151 treatment (tumor + CRT, F(1,27)=8.0, p<0.01; tumor + CRT x time, F(2, 54)=14.3, p<0.001; H-151, F(1, 27)=0.02, not significant) (**Fig. 5D**).

## 4. Discussion

The present findings show that cancer and CRT activate the cGAS-STING pathway. This activation positively contributes to cancer-related fatigue, measured by decreased voluntary wheel running activity in mice.

Activation of the cGAS-STING pathway has mainly been studied in cancer immunology because of its pivotal role in the progression and dissemination of tumors. This pathway can be engaged by tumor cells’ intrinsic mechanisms and by non-autonomous mechanisms occurring in immune and stromal cells in the tumor (26). Depending on the context, the outcome can be an inhibition or, conversely, an augmentation of tumor progression. Cancer therapy, including chemotherapy and irradiation, can also directly engage the cGAS-STING pathway, which usually promotes tumor eradication due to its immunogenic potential. This explains the current and growing focus on identifying means to harness cGAS-STING in a controlled manner to potentiate the effects of current anti-cancer treatments.

The tumor-centric nature of research on the cGAS-STING pathway in cancer has led to the neglect of several other aspects of this pathway, including the possibility of its engagement in different parts of the body beyond the tumor, and the mechanisms underlying its tumor-independent effects. As already demonstrated in cancer metabolism syndrome, the soluble mediators that are released by tumor cells can act at a distance from their site of production either because they are transported as such or included in microvesicles that are released in the general circulation, or because they activate locally the sensory nerves that innervate the tumor and form a tumor-brain circuit (27, 28). Our findings here support this possibility. Although we utilized tumor cells that do not express cGAS-STING, there is still the possibility that self-DNA released by tumor cells or non-tumor cells in the harsh tumor microenvironment engages cGAS-STING. Our results on the expression of the cGAS-STING pathway in the tumor attest to this possibility. Although we did not study how activation of the cGAS-STING pathway propagates from the tumor to distant organs such as the liver, it can be proposed that what happens in this organ is the result of the cellular stress imposed by the tumor on liver metabolism to meet its energy requirements (29). Local engagement of the cGAS-STING pathway could explain why inflammation develops in the liver and other organs recruited by cancer metabolism syndrome. To understand the way this cascade of events ultimately develops, it would still be necessary to study them at different time points during the progression of the tumor and to intervene on specific cell types to delete elements of the cGAS-STING pathway.

The objective of the present study was more focused. Given the limited knowledge of the extra-tumoral consequences of activation of the cGAS-STING pathway in response to tumor progression and to cancer therapy, we decided to focus on the possible involvement of this pathway in cancer-related fatigue. This choice was additionally motivated by the emphasis put on inflammation as the main causal factor by those who study fatigue and other cancer-related symptoms (7). We already know that various DAMPs produced by tumor cells can activate inflammation locally and at a distance from the tumor. However, it remains that “aberrant DNA recognition via the cGAS–STING pathway is of particular relevance for endogenous sensing of transforming or transformed cells, both at basal levels and following cancer treatment” (26). This means that the possible involvement of this pathway in the pathophysiology of cancer-related fatigue cannot be ignored. Recently, transcriptional signatures of type I interferon signaling were shown to correlate with fatigue severity in breast cancer patients receiving chemotherapy, radiotherapy, or CRT (30). The cellular sources of this type I interferon signature and the signaling pathways that are responsible for its occurrence in peripheral blood mononuclear cells of breast cancer patients have not yet been identified. However, as the cGAS-STING pathway is the major innate immune network activated by CRT and the endogenous nucleic acids released from dying tumor cells and injured tissues, it is highly plausible that cGAS-STING lies upstream of the fatigue-associated interferon signature.

In the present study, we investigated the role played by the activation of the cGAS-STING pathway in cancer using association and intervention studies. Both approaches converged to show that this pathway qualifies as a candidate target for minimizing fatigue associated with cancer and CRT. This does not necessarily mean that the cGAS-STING pathway is a feasible target for treating fatigue or other cancer treatment-related symptoms. Given the role of the STING pathway in modulating the tumor immune microenvironment, inhibiting it to control fatigue may be counter-productive for the treatment of cancer. However, a growing body of evidence suggests that targeted intervention in STING signaling or downstream factors can limit some of the off-target effects of chemotherapy and/or radiotherapy (e.g., cardiotoxicity (31)(12); acute kidney injury (32); hearing loss (33). Still, there are several important limitations of this work. Notably, our study was only conducted in only one model of cancer, and we did not address the possible existence of sex differences in the engagement of the cGAS-STING pathway. Therefore, the present findings must be viewed as supporting evidence for the potential role of this pathway in fatigue and not definitive. Future studies involving additional tumor models and time course experiments will be helpful in clarifying the sequence of events leading to cGAS-STING activation, which may reveal a unique window for intervention in cancer- and treatment-related fatigue.

## Acknowledgments

Supported by NIH-NCI R01 CA193522 and an MD Anderson Cancer Center Support Grant (P30 CA016672). A. Phillip West was supported by The Jackson Laboratory Cancer Center Support Grant (P30 CA034196). Abate Bashaw was supported by a Brooks Scholar Award from The Jackson Laboratory Cancer Center.

## CRediT authorship contribution statement

**Brandon Chelette:** Investigation, Visualization, Data curation, Formal analysis, Writing & editing. **Abate Bashaw:** Investigation, Visualization, Data curating, Formal analysis, Writing & editing. **Joshua D. Bryant:** Data curation, Formal analysis, Visualization. **Kiersten S. Scott:** Formal analysis, Investigation. **Chinenye Chidomere**: Investigation, Visualization, Data curating. **A. Phillip West:** Conceptualization, Methodology, Supervision, Formal analysis, Writing – original draft, review & editing. **Robert Dantzer:** Conceptualization, Formal analysis, Funding acquisition, Methodology, Project administration, Resources, Supervision, Validation, Visualization, Writing – original draft, review & editing.

**Supplementary Table 1.**
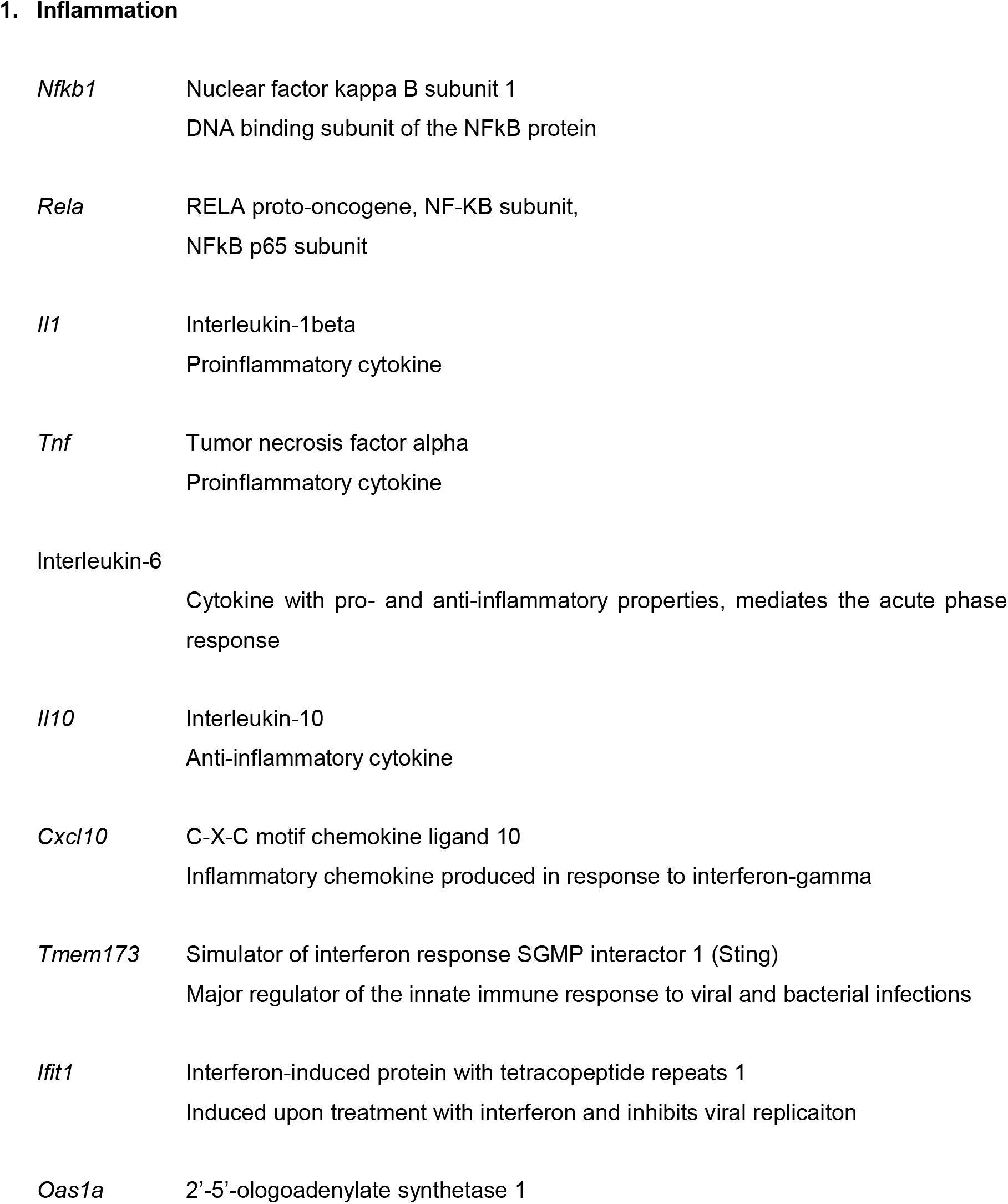
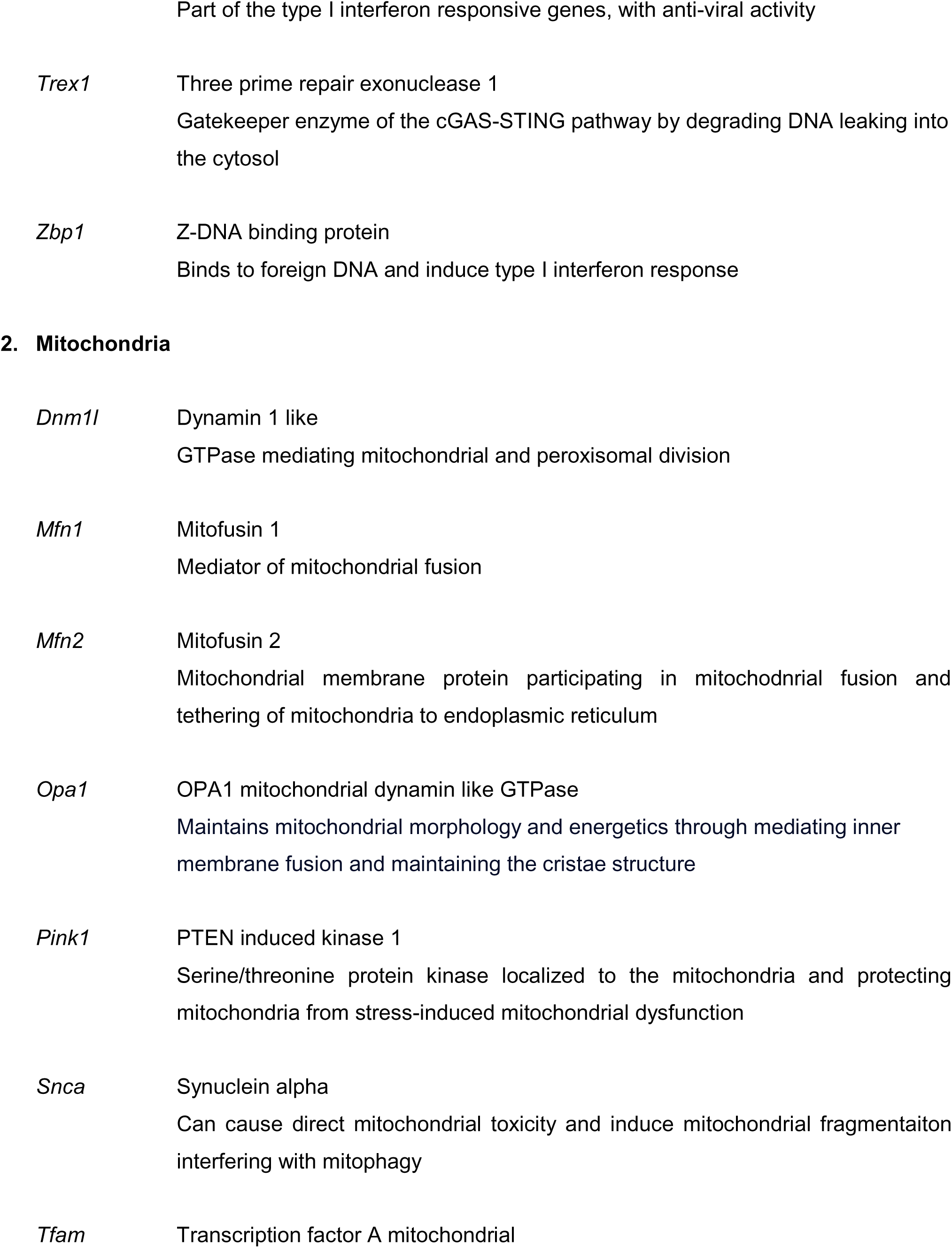
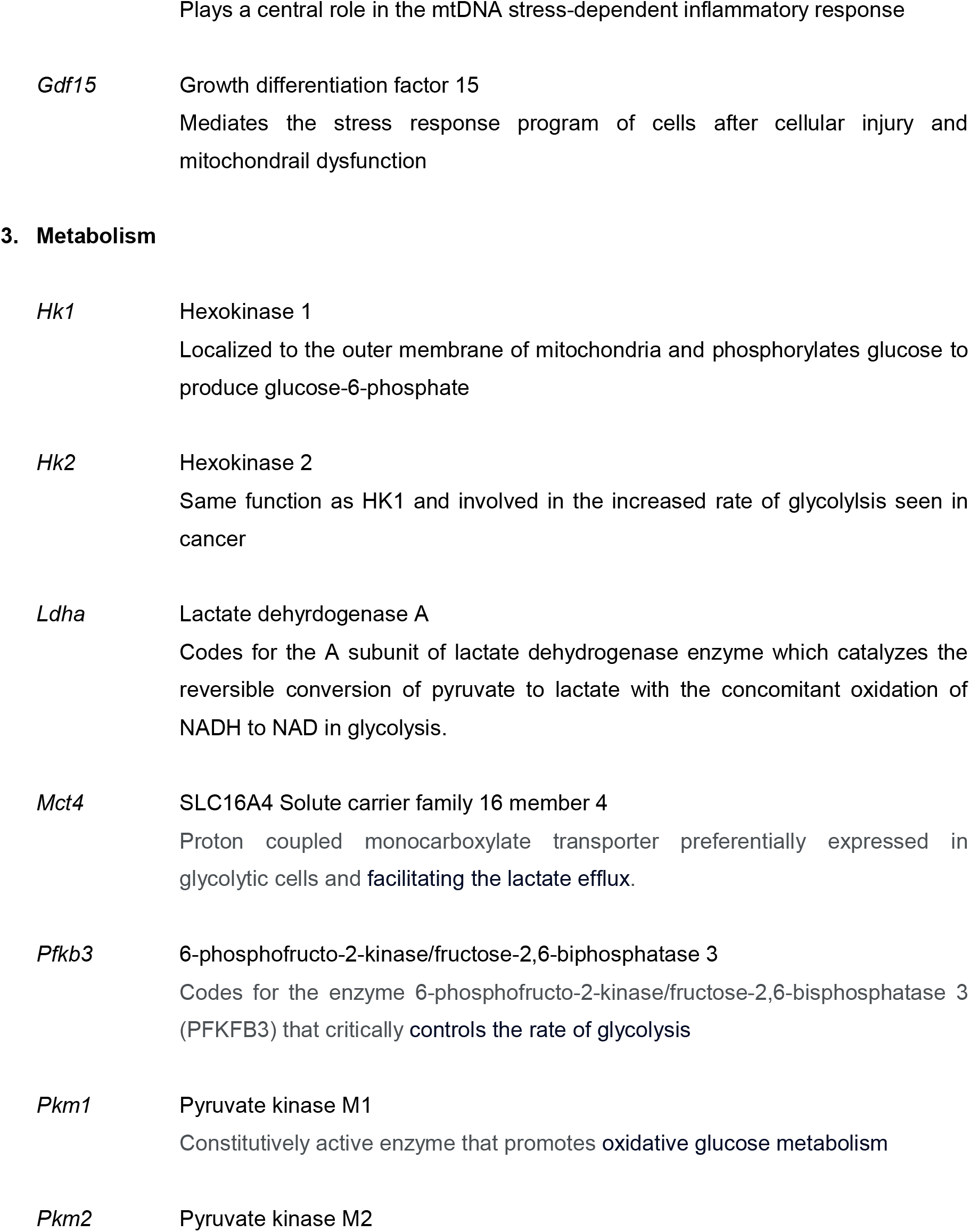
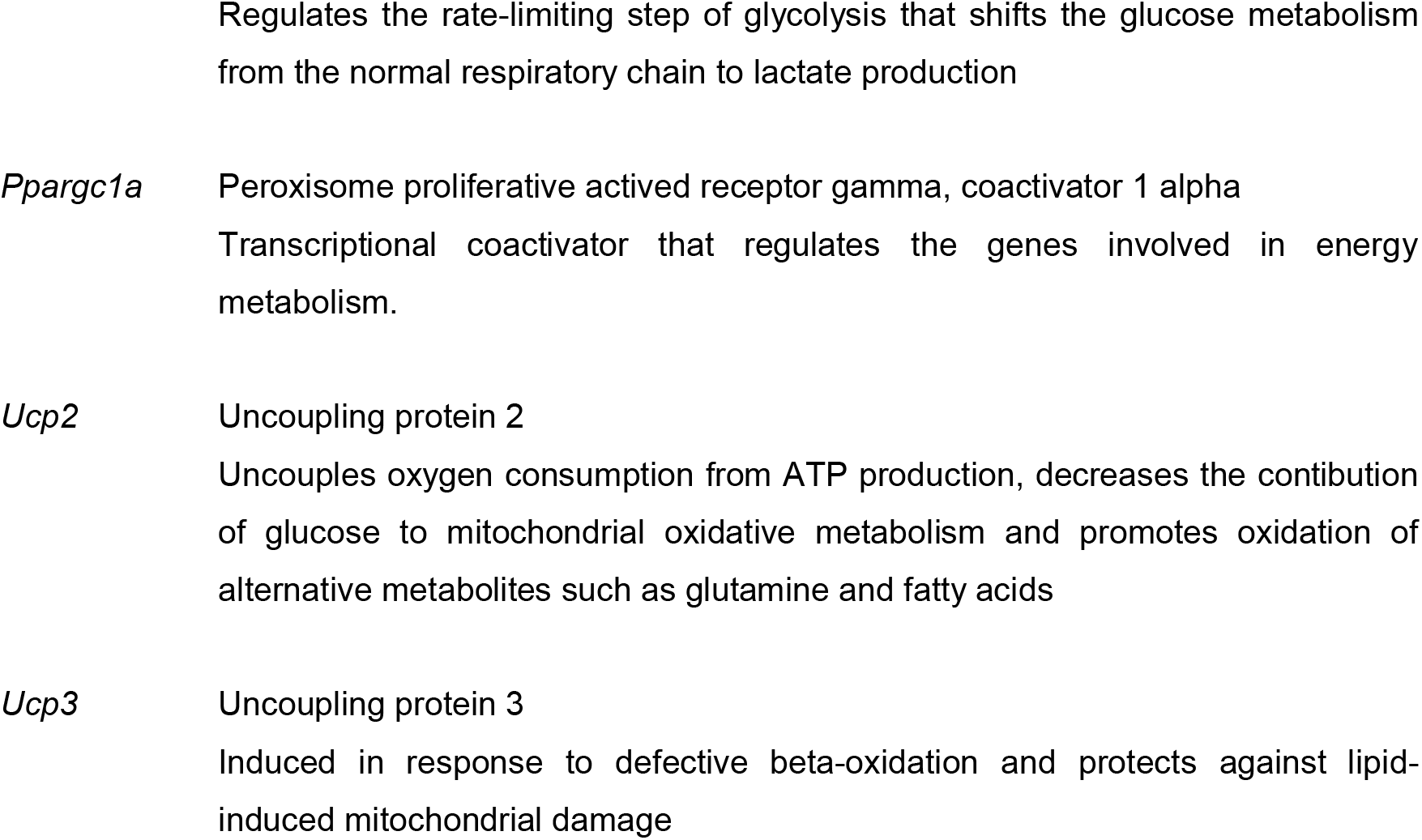
List of select genes (symbol followed by description and function of the coded protein)

